# Molecular specification of cortico-brainstem versus corticospinal projection neurons in development

**DOI:** 10.1101/2022.05.31.494253

**Authors:** Julia Kaiser, Payal Patel, Friederike Dündar, Jimena Perez-Tetuan, Nirupama Angira, Eytan Sieger, Vibhu Sahni

## Abstract

Skilled motor control requires precise connections between subcerebral projection neurons (SCPN) in the cerebral cortex and their appropriate subcerebral targets in the brainstem or spinal cord. The brainstem is an important motor control center and cortical projections to the brainstem serve distinct motor control functions than corticospinal projections. However, mechanisms controlling cortico-brainstem versus corticospinal projections during development remain unknown. Here, we show that the transition between the brainstem and cervical cord distinguishes cortico-brainstem from corticospinal neurons from the earliest stages of SCPN axon extension in white matter. We used high throughput single-cell RNA sequencing of FACS-purified SCPN, retrogradely labeled from either the cerebral peduncle (labeling both cortico-brainstem and corticospinal neurons) or the cervical cord (labeling corticospinal neurons only) at critical times of axon extension. We identify that cortico-brainstem and corticospinal neurons are molecularly distinct: We establish *Neuropeptide Y* (*Npy*) as specifically enriched in cortico-brainstem neurons in lateral cortex, while *CART prepropeptide* (*Cartpt*) delineates cervical-projecting corticospinal neurons. Our results highlight molecular specification of cortico-brainstem vs. corticospinal projections well before these axons reach their appropriate segmental target and suggest a broad molecular program over SCPN axon targeting to distinct subcerebral targets early in development. These findings are likely to inform future investigations of motor circuit development, as well as approaches aimed at enhancing motor recovery after central nervous system damage.

**Highlights:**

- Cortico-brainstem neurons (CBN) limit their axon extension to supraspinal levels from the earliest time points of white matter axon extension in development.
- CBN can be molecularly delineated from corticospinal neurons (CSN) even at these initial time points.
- Molecular diversification of developing subcerebral projection neurons occurs across at least two axes: cortical location (medial vs. lateral) and projection targeting specificity (brainstem vs. spinal)
- Within lateral cortex, Neuropeptide Y (*Npy*) is expressed by CBN, while CART prepropeptide (*Cartpt*) expression delineates cervical-projecting CSN.

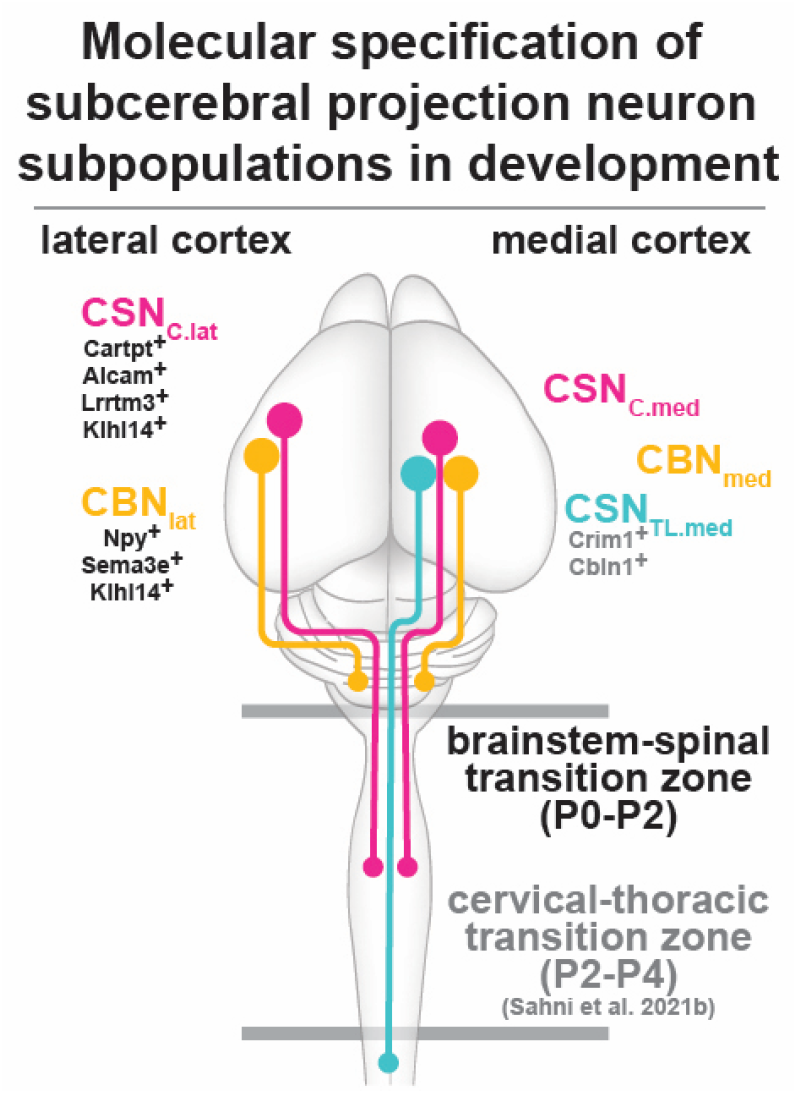

## Introduction

For skilled motor control, neocortical subcerebral projection neurons (SCPN) must make precise connections with subcerebral targets in the brainstem and spinal cord (Lemon, 2008; Levine et al., 2012; Sahni et al., 2020). This segmentally appropriate connectivity is essential for distinct functions in motor control. Historically, cortico-brainstem neurons (sometimes referred to as “cortico-bulbar” neurons) are known to control facial movement, while corticospinal neurons are thought to be responsible for arm, trunk, and leg motor control (Schieber, 2007). The brainstem, however, has recently emerged as a critical motor control center that is also responsible for skilled limb movements (Esposito et al., 2014; Ruder et al., 2021), and receives significant cortical projections (Economo et al., 2016, 2018). In this regard, recent work has further identified that cortico-brainstem projections provide specific modulatory roles required for dexterous movement that are distinct from corticospinal projections (Conner et al., 2021; Moreno-Lopez et al., 2021). Further, both corticospinal projections, as well as cortico-brainstem and brainstem-spinal projections show a high degree of plasticity after central nervous system (CNS) damage that contributes to motor recovery (Bachmann et al., 2014; Kaiser et al., 2019), with these distinct projections potentially mediating distinct elements of functional motor recovery. However, the mechanisms that specify these anatomically and functionally distinct cortical projections during development remain largely unknown.

Prior work from several groups has identified transcriptional regulators that not only distinguish SCPN from other neocortical projection neuron types e.g. corticothalamic projection neurons or callosal projection neurons (Fame et al., 2011; Franco & Müller, 2013; Greig et al., 2013; Leone et al., 2008; Lodato et al., 2015; Molyneaux et al., 2007), but also control critical aspects of their specification, differentiation, and subcerebral axonal connectivity (Arlotta et al., 2005; Cederquist et al., 2013; B. Chen et al., 2005; J. G. Chen et al., 2005; Greig et al., 2016; Han et al., 2011; Joshi et al., 2008; Kwan et al., 2008; Lai et al., 2008; Leone et al., 2015; Lodato et al., 2014; McKenna et al., 2011, 2015; Molyneaux et al., 2005; Özdinler & Macklis, 2006; Shim et al., 2012; Tomassy et al., 2010; Woodworth et al., 2012). Our recent work identified that corticospinal neurons (CSN) exhibit striking axon targeting specificity at the transition between cervical and thoracic spinal segments (Sahni, Shnider, et al., 2021). This axon extension specificity occurs at the level of the spinal white matter prior to axonal collateralization or synapse formation and is controlled by CSN-intrinsic determinants. CSN that extend axons proximal to this transition zone are molecularly distinct from CSN that extend axons beyond it and genes differentially expressed between these CSN subpopulations control their differential axon targeting (Sahni, Itoh, et al., 2021).

In this report, we establish that the transition between the brainstem and the spinal cord represents an additional, more rostral site for differential SCPN axon targeting specificity in early development. This developmental specificity distinguishes cortico-brainstem neurons, which do not extend axons past this site, from corticospinal neurons. We further establish that these distinct SCPN subsets can be molecularly delineated even before their differential axon targeting is fully established, indicating early molecular specification of cortico-brainstem versus corticospinal projection neurons.

## Results

### Cortico-brainstem neurons limit their axon extension to supraspinal levels from initial stages of SCPN axon extension

Subcerebral projection neurons (SCPN) extend axons from their neocortical location in layer V to subcerebral targets throughout the rostro-caudal extent of the neuraxis, ultimately innervating either brainstem (cortico-brainstem projection neurons, CBN) or spinal (corticospinal projection neurons, CSN) targets (Fig 1A). SCPN axon extension specificity at the transition between cervical and thoracic spinal segments is established from the earliest stages of axon extension into the spinal cord, distinguishing bulbar-cervical from thoraco-lumbar projecting neurons (Sahni, Shnider, et al., 2021). This suggested that there are additional points throughout the neuraxis at which similar axon extension specificity might occur. We therefore analyzed whether CBN axon extension is restricted to supraspinal levels from the earliest times of axon elongation, well before collateral branching or innervation occurs. To address this question, we retrogradely labeled SCPN from the white matter at two distinct levels of the neuraxis: we first injected a retrograde tracer into the cerebral peduncles at P0, labeling all SCPN (both CBN and CSN) (Fig.1 A). In a separate set of mice, we injected a retrograde tracer into the dorsal funiculus of the cervical spinal cord at P2, labeling CSN only (Fig. 1B). Labeled SCPN (CBN + CSN) are broadly distributed in layer V throughout the rostro-caudal extent of sensorimotor cortex (Fig 1C). In contrast, retrograde tracing of CSN labels only a subset of all SCPN (Fig 1D). This indicates that a significant number of SCPN axons extend to the brainstem but never enter the cervical spinal cord, suggesting that a majority of developing SCPN limit their axon extension to supraspinal levels (i.e., CBN).

**Fig 1.**
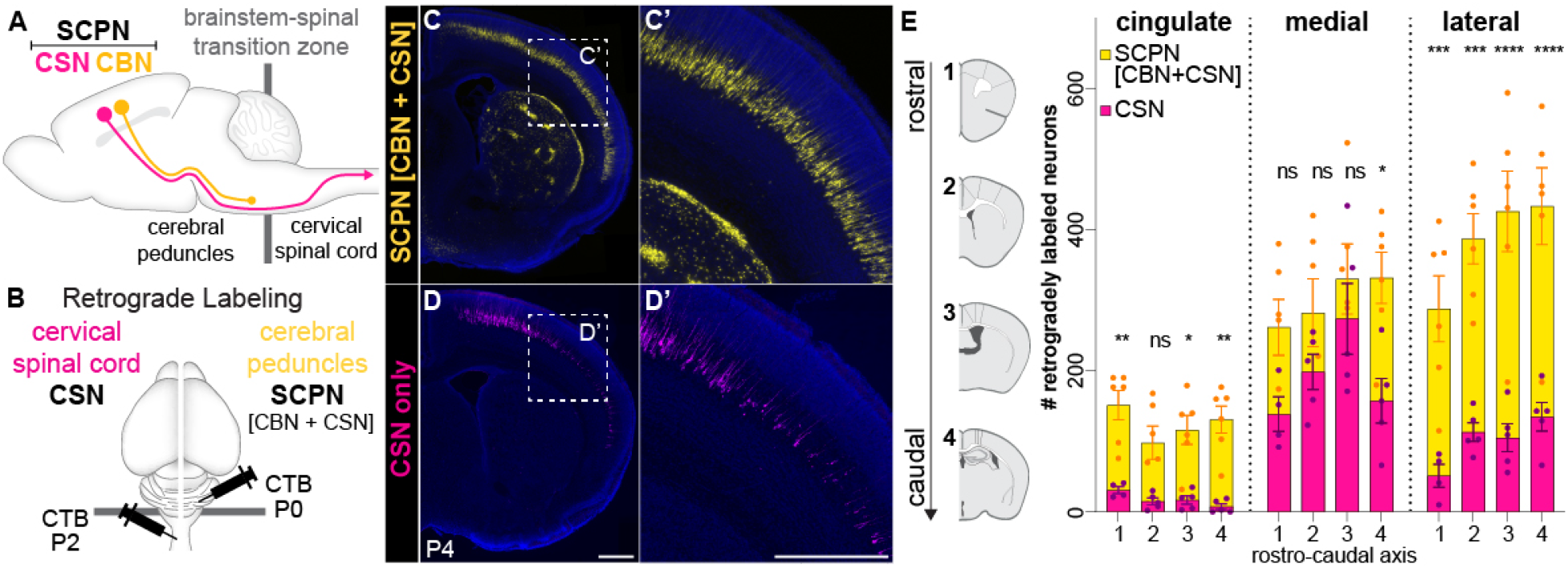
The brainstem-spinal transition zone is established early in development. (A) Subcerebral projection neurons extend axons to subcerebral targets. This broader population comprises both cortico-brainstem neurons (CBN, yellow) that project only to the brainstem, as well as corticospinal neurons (CSN, magenta) that extend to the spinal cord. (B) Experimental scheme: Retrograde labeling from cerebral peduncles labels all SCPN (CBN +CSN) while retrograde labeling from the cervical spinal cord only labels CSN. Injections were done at the time point of initial axon extension– P0 from the cerebral peduncles; P2 from cervical spinal cord. (C) Retrogradely labeled SCPN reside throughout the medio-lateral extent of sensorimotor cortex. (D) The majority of labeled CSN reside in medial sensorimotor cortex, with few CSN within the lateral sensorimotor cortex (zoom-in views), (E) Quantification of retrogradely labeled neurons. Almost all SCPN in cingulate cortex are CBN; the majority of CSN reside in medial sensorimotor cortex together with CBN, only a minority of SCPN in lateral sensorimotor cortex are CSN. n = 6 (SCPN), n = 5 (CSN). Scale bar, 500µm. *P < 0.05, **P < 0.01, ***P < 0.001.

Quantification of retrogradely labeled neurons in the two experiments shows that the distribution of CBN vs. CSN varies across sensorimotor cortex (Fig. 1E). The vast majority of CSN reside in medial sensorimotor cortex (this includes primary motor cortex), intermingled with CBN. The cingulate cortex is almost completely devoid of CSN; almost all SCPN in cingulate cortex are CBN. The lateral sensorimotor cortex is similarly enriched for CBN. Although CSN represent a minority of the overall SCPN in lateral sensorimotor cortex, they are evenly distributed throughout the rostro-caudal extent of lateral cortex (Fig 1E). We additionally validated these findings via dual retrograde labeling of both CSN and SCPN in the same mouse and find identical results (Supplemental Fig 1). Taken together, these results highlight that CBN axons limit their axon extension to supraspinal levels from the earliest stages of axon elongation. This establishes a brainstem-spinal transition zone as an additional site where SCPN axons exhibit differential axon extension specificity early in development.

### Molecular specification of subsets of CBN and CSN

The striking axon extension specificity by CBN and CSN suggested that these subpopulations are molecularly specified for their differential axon extension early in development. Thus, we FACS-purified retrogradely labeled SCPN (labeled from the cerebral peduncles) versus CSN (labeled from the cervical cord) and performed differential gene expression analysis using single-cell transcriptomics. Integration of SCPN and CSN samples allowed us to transcriptionally delineate CBN, as they were present only in SCPN but not CSN samples. Samples were collected at P1 and P3, representing critical developmental times of white matter axon extension: P1 is prior to, and P3 is immediately after CSN axon extension into the spinal cord. By integrating datasets across these critical developmental times, we analyzed temporal dynamics of gene expression that correlate with the dynamics of differential axon targeting (Fig 2B). In total, neurons had a median of 3498 unique molecular identifiers (UMIs), a median of 1904 detected genes per cell, and a median of 6.98 % of mitochondrial genes. We used a published single-cell dataset of P0 mouse cortex (Loo et al., 2019) to annotate distinct cell types in a semi-automated manner. We find some contamination of our samples by other cell types, however, FACS purification enabled significant enrichment of layer V-VI neurons (56.7%, Fig 2B, dashed box) that were used for downstream analysis. Unbiased clustering of these layer V-VI neurons identified transcriptionally distinct clusters. We hypothesized that axonal extension diversity would, at least partially, underlie these transcriptional differences.

**Fig 2.**
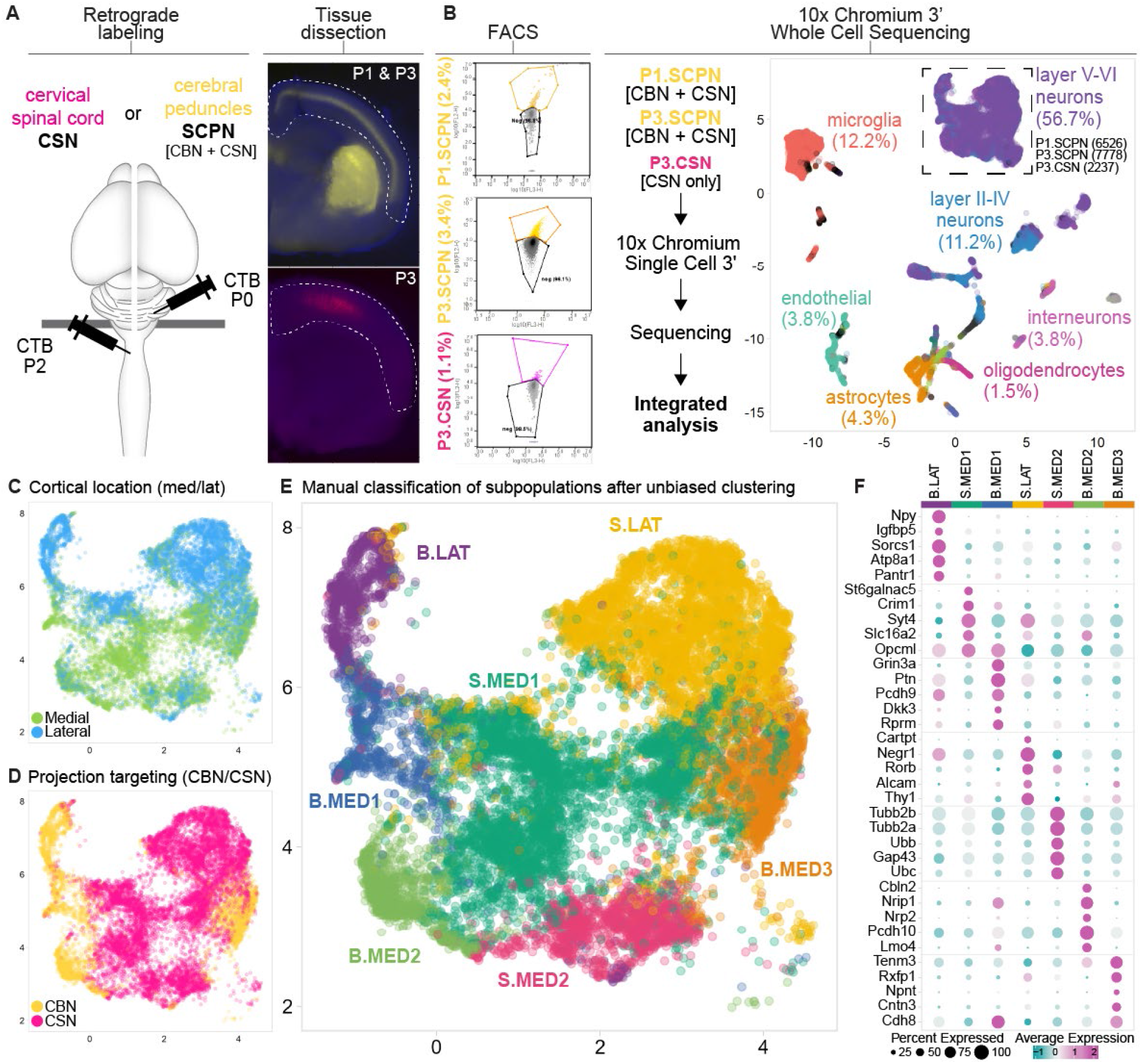
Molecular specification of SCPN subpopulations based on projection targeting specificity and cortical location. (A) Experimental Scheme: SCPN or CSN were retrogradely labeled (SCPN at P0, CSN at P2), cortices were dissociated at P1 (SCPN) or P3 (SCPN or CSN) and whole cell suspensions were FACS purified. (B) Sequencing based off of the 10x Genomics platform allowed for an integrated analysis of all 3 samples (P1.SCPN, P3.SCPN, P3.CSN). We find some contamination by non-neuronal cell types, but highly enriched for layer V-VI neurons (annotation based on P0 cortical cell type data (Loo et al., 2019)). (C) SCPN cortical location in medial (“MED”) vs. lateral (“LAT”) cortex (Sahni, Shnider, et al., 2021). (D) Projection targeting specificity: clusters that had cells only from SCPN samples were classified as “CBN” (brainstem-projecting, “B”), while clusters that contained cells from both SCPN and CSN samples were classified as “CSN” (spinal-projecting, “S”). (E) Manual classification of clusters identified using unbiased clustering (k50) on data from both cortical location and projection targeting specificity identified seven subclusters that we further characterized. (F) Heatmap of top 5 differentially expressed genes between the 7 clusters. Npy and Cartpt are the top differentially expressed genes for B.LAT and S.LAT, respectively.

Our prior work identified that SCPN in lateral vs. medial sensorimotor cortex extend segmentally distinct projections – SCPN in lateral cortex extend projections exclusively to brainstem and cervical cord (bulbar-cervical), while SCPN in medial cortex are more heterogeneous with distinct subsets projecting to either bulbar-cervical or thoraco-lumbar segments (Sahni, Shnider, et al., 2021). To investigate whether molecular delineation within SCPN can be attributed to cortical location, we used transcriptional profiling data from this previous study, which identified differentially expressed genes between medial vs. lateral SCPN, to annotate clusters by medial vs. lateral cortical location (Fig 2C). We further annotated projection targeting specificity based on sample subtype (i.e., clusters that were only present in SCPN but not CSN samples were annotated as “CBN”, while clusters that were present in both samples were annotated as “CSN”, Fig 2D). Using these annotations, we manually annotated the transcriptionally distinct clusters based on their 1) projection targeting specificity (cortico-brainstem “B” vs. corticospinal “S”); and 2) cortical location (medial (“MED”) vs. lateral (“LAT”)) (Fig 2E).

Consistent with previous anatomical data (Sahni, Shnider, et al., 2021), we find that medial cortex SCPN are represented by more transcriptionally distinct clusters (5 out of 7 total) than SCPN in lateral cortex. This potentially represents the greater diversity of subcerebral projection targets within medial SCPN. The medial cortex SCPN clustered into 5 distinct subclusters – 2 corticospinal clusters (S.MED1, 2), and 3 cortico-brainstem clusters (B.MED1, 2, 3). In contrast, lateral cortex SCPN segregated into 2 clusters – specifically one cluster each for CBN (“B.LAT”), and CSN (“S.LAT”) (purple and yellow clusters, Fig 2E), corroborating our previous findings that subcerebral projections arising from lateral cortex are relatively more homogeneous than projections from medial cortex. Lateral SCPN axons extend only to the brainstem and cervical spinal cord (Sahni, Shnider, et al., 2021), thus, the spinal lateral cluster (“S.LAT”) should be specific to cervical projecting CSN (CSN_C-lat_).

We next identified the top differentially expressed genes between all clusters (Fig 2F). Genes identified as differentially expressed between all clusters further corroborated our annotations (Fig 2F). Genes previously identified as specific to thoraco-lumbar-projecting CSN (CSN_TL_), e.g., Crim1 (Sahni, Itoh, et al., 2021), delineate the medial corticospinal cluster “S.MED1” (Fig. 2F), suggesting that “S.MED1” represents CSN_TL_. Since the lateral cortex has 2 well-defined clusters that were annotated as cortico-brainstem vs. corticospinal neurons, we next selected genes that were differentially expressed between these subpopulations for more detailed expression and projection analyses. For the two lateral clusters, “B.LAT” and “S.LAT”, the top differentially expressed genes are *Neuropeptide Y (Npy)* and *CART prepropeptide* (*Cartpt)*, respectively (Fig 2F).

### *Cartpt* is enriched in corticospinal neurons in lateral cortex and its expression increases as corticospinal axons extend into the spinal cord

Analysis of the top differentially expressed genes highlights *Cartpt* as enriched in CSN_C-lat_ (“S.LAT”, Fig 2F). *Cartpt* encodes a preproprotein that is cleaved into multiple distinct neuroactive CART peptides that are widely distributed in the CNS and involved in regulating many processes, including food intake and the maintenance of body weight, reward, and endocrine functions (Rogge et al., 2008). Interestingly, we find that *Cartpt* expression increases from P1 to P3, as corticospinal axons are extending into the spinal cord (Fig 3A). To validate these single-cell RNA-seq data, we used RNAScope (*in situ* hybridization) to investigate *Cartpt* expression within the cortex at P3. We find that *Cartpt* is specifically expressed in lateral but not medial cortex (Fig. 3C-D). We also combined this expression analysis with retrograde labeling from the cervical spinal cord to label corticospinal neurons and investigated whether *Cartpt* is expressed by CSN in lateral cortex. *Cartpt* is expressed by almost all retrogradely labeled CSN in lateral cortex (Fig 3C, Supplemental Fig 2). This confirms the transcriptomic profiling data and suggests that *Cartpt* is specifically expressed by cervical projecting corticospinal neurons at the time when CSN axons extend into the spinal cord. We also investigated the expression of other top differentially expressed genes in the “S.LAT” cluster: *Alcam* and *Lrrtm3*. Using similar expression analyses of combining RNAScope with retrograde labeling, we find that *Alcam* and *Lrrtm3* are also expressed by corticospinal neurons in lateral cortex, thus providing additional molecular candidates that delineate CSN_C-lat_ from CBN (Supplemental Fig 2). To our knowledge, these genes provide the first developmental identifiers of cervical-projecting CSN during the initial stage of corticospinal axon extension into the spinal cord.

**Fig 3.**
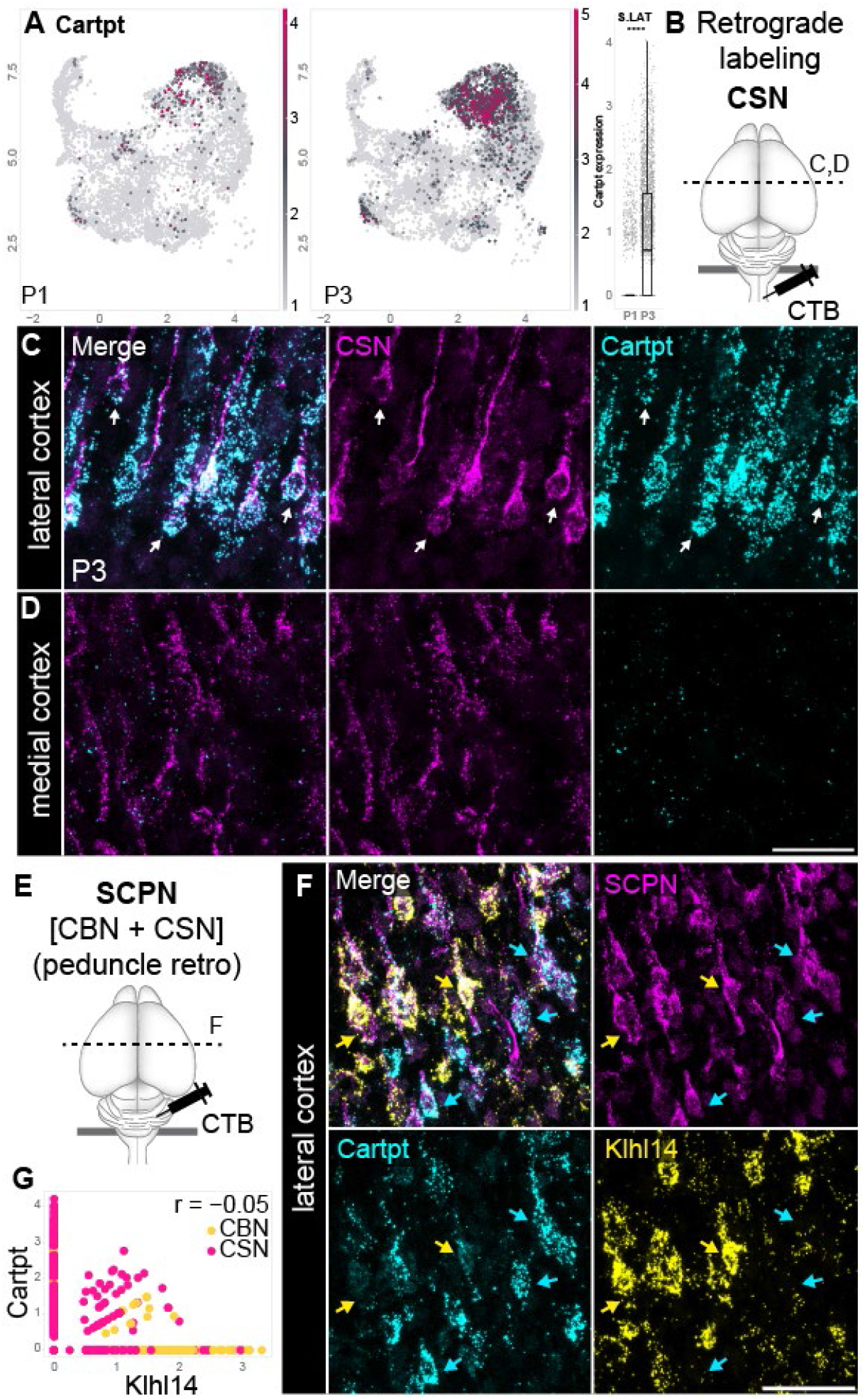
Cartpt expression is specific to corticospinal neurons in lateral cortex. (A)UMAP of Cartpt expression shows specific expression within cluster “S.LAT”, with an enrichment for Cartpt at P3. (B) CSN were retrogradely labeled at P2, and Cartpt expression in the cortex analyzed using RNAScope at P3. (C) P3 brain showing that in lateral cortex Cartpt is expressed by retrogradely labeled CSN (white arrows). (D) Cartpt is not expressed in medial cortex. (E) All SCPN were retrogradely labeled from the cerebral peduncle at P0. (F) P3 brain showing that in lateral cortex, both Cartpt and Klhl14 are expressed by retrogradely labeled SCPN; however, Cartpt and Klhl14 do not colocalize, indicating that Cartpt^+^ SCPN (cyan arrows) are molecularly distinct from Klhl14^+^ SCPN (yellow arrows). (G) In the 10x dataset, there is little correlation between Cartpt and Klhl14. Furthermore, while almost all Cartpt expression occurs in CSN, Klhl14 is expressed by both CSN and CBN, with high-level Klhl14^+^ expression occurring in CBN.

Our previous work identified *Kelch-like 14 (Klhl14)* as specifically expressed by bulbar-cervical SCPN in lateral cortex (Sahni, Shnider, et al., 2021). We therefore investigated whether *Cartpt*^+^ SCPN in lateral cortex are also *Klhl14*^+^ or whether they represent a distinct subpopulation (Fig 3E-G). We retrogradely labeled all SCPN and performed RNAScope for both genes. While both genes are expressed by SCPN (Fig 3E), there is no overlap between *Cartpt* and *Klhl14* (Fig 3F,G), indicating that they are expressed by distinct SCPN subpopulations in lateral cortex. Correlation analysis of cellular expression in the single-cell RNA-Seq data confirms these findings (Fig. 3G). While *Cartpt* is almost exclusively expressed by CSN in lateral cortex, Klhl14 appears to be expressed by both CBN and CSN (Fig 3G). This indicates that Klhl14 is expressed by a broader population of both cervical and brainstem projecting SCPN (“bulbar-cervical”) in lateral cortex, as was previously reported (Sahni, Shnider, et al., 2021).

### Npy expression delineates cortico-brainstem neurons in lateral cortex

The most specifically expressed gene for the brainstem lateral cluster (“B.LAT”) is *Npy* (Fig 2F), which is particularly enriched in this cluster and shows higher expression at P1 than P3 (Fig 4A). *Npy* is expressed throughout the central, peripheral, and autonomic nervous systems and controls multiple aspects of normal physiology. Until recently, *Npy* was known to be expressed by cortical interneurons (Karagiannis et al., 2009); however, consistent with our findings, recent evidence indicates that *Npy* is also expressed by a subset of excitatory projection neurons during early cortical development (Di Bella et al., 2021). Using RNAScope, we confirmed that *Npy* expression co-localizes with retrogradely labeled SCPN (Fig 4C) in lateral cortex. While *Npy* is expressed in medial cortex, this expression does not overlap with any retrogradely labeled SCPN, indicating that this expression is likely by other neocortical neuron types (Fig 4D).

**Fig 4.**
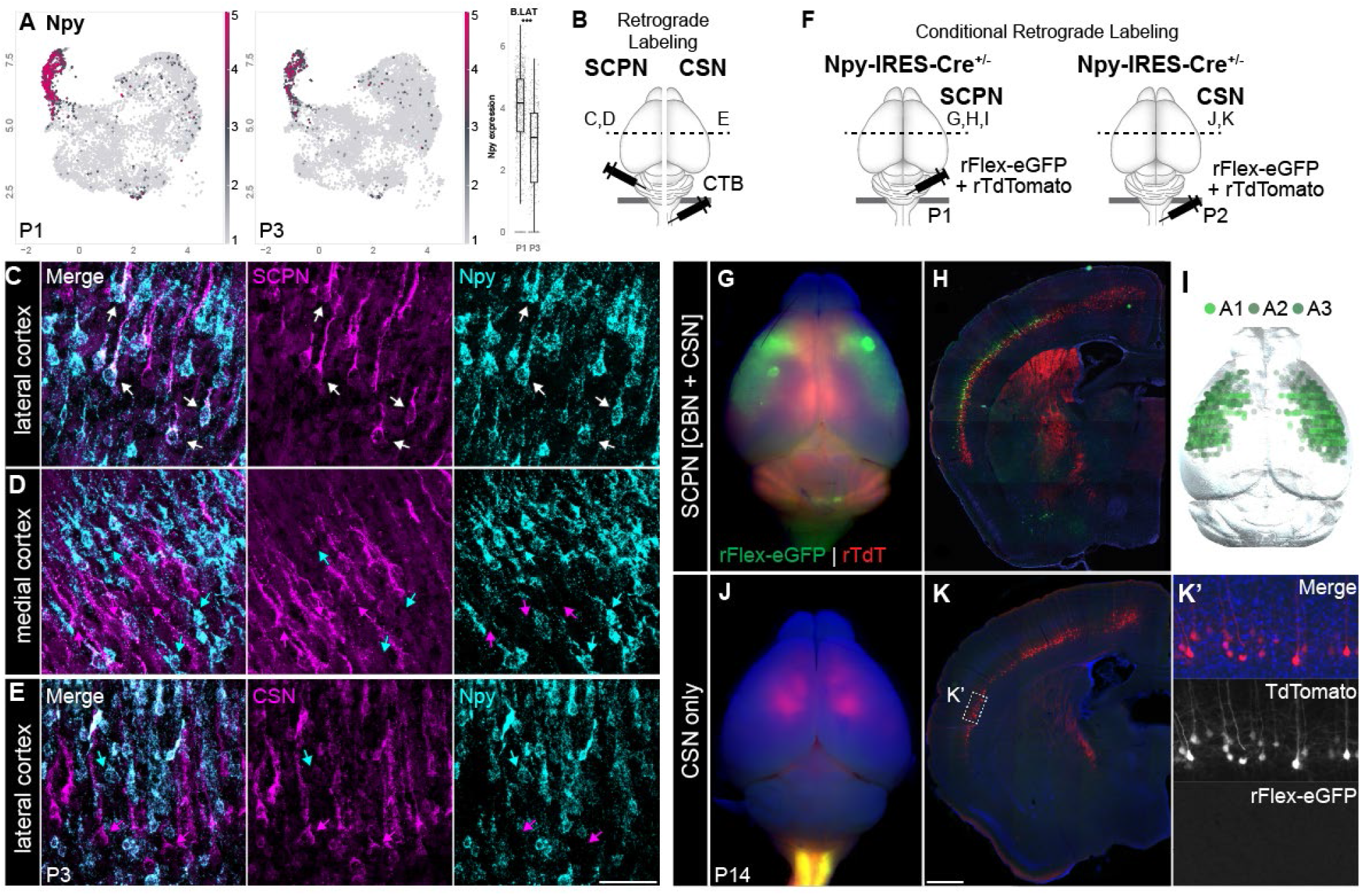
Npy is specifically expressed by CBN and not CSN in lateral cortex. (A) UMAP of Npy expression demonstrates the specificity to cluster “B.LAT” with higher enrichment at P1 than P3. (B) Schematic representation of retrograde labeling for CSN (retrograde tracing from cervical spinal cord at P2) and SCPN (peduncle retrograde tracing at P0). (C,D) RNAScope for Npy in lateral cortex (C) or medial cortex (D) with SCPN retrograde labeling at P3. (E) RNAScope for Npy with CSN retrograde labeling. (F) Conditional viral labeling of Npy^+^ projection neurons: In Npy- IRES-Cre knock-in reporter mice, rFlex-GFP was injected into the cerebral peduncles at P1 (left) to label Npy^+^ SCPN or the spinal cord at P2 (right) to label Npy^+^ CSN. rTdTomato was co-injected as a control to label all SCPN/CSN, respectively. Mice were perfused at P14. (G-I) Conditionally labeled Npy^+^ projection neurons are located in lateral cortex as seen in the whole mount (G), coronal section of the same brain (H) as well as 3D representations from 3 distinct mice (I). (J). While TdTomato^+^ retrogradely labeled CSN can be seen throughout the cortex, we do not see conditionally labeled Npy^+^ projection neurons in neither the whole mount overview (J), nor coronal sections (K,K’’) confirming that Npy^+^ SCPN are not CSN.

We next investigated *Npy* specificity between CBN and CSN. We again performed retrograde labeling from the cervical cord to label CSN and analyzed whether *Npy*^+^ SCPN colocalize with labeled CSN in lateral cortex. We find no overlap between *Npy* expression and retrogradely labeled CSN (Fig 4E), which strongly suggests that *Npy* is specifically expressed by CBN and not CSN in lateral cortex.

### Npy^+^ SCPN extend axons to the brainstem but not the spinal cord

To further corroborate our RNAScope finding that *Npy* is specific to CBN and excluded from CSN, we performed conditional retrograde labeling using retro-AAVs in Npy-IRES-Cre knock-in reporter mice (Milstein et al., 2015). As in the previous experiments, we retrogradely labeled SCPN (both CBN and CSN) by injecting a control retro-TdTomato along with a Cre-dependent retro-Flex-eGFP into the cerebral peduncles (Fig 4F, left schematic). We first validated the location of eGFP- labeled (*Npy*^+^) SCPN: while control TdTomato^+^ SCPN are present in both medial and lateral cortex, eGFP labeled *Npy*^+^ SCPN are indeed located predominantly in lateral cortex (Fig 4G-I). This further confirms the results from our single-cell RNA-seq data regarding cortical location, as well as our previous RNAScope analyses.

We next investigated whether *Npy* expression is indeed specific to CBN and not CSN. For this, we performed retrograde labeling from the cervical spinal cord using the same conditional labeling strategy (Fig 4F, right schematic). If *Npy*^+^ SCPN are exclusively CBN, we would not expect any eGFP-labeled (*Npy*^+^) CSN using this labeling approach. Indeed, we find that while control retro-TdTomato successfully labels CSN in both medial and lateral sensorimotor cortex (Fig 4J,K), we do not find eGFP-labeled (*Npy*^+^) CSN when traced from the spinal cord (Fig 4K,K’’). This further confirms that *Npy* expression within SCPN is specific to CBN in lateral cortex (B.LAT) during development. Together with the earlier *Cartpt* expression analyses, these results further validate the single-cell sequencing data and highlight that projection targeting diversity is a major axis of transcriptional diversity within SCPN, with *Cartpt* specifically delineating CSN_C-lat_ and *Npy* specifically expressed by CBN.

## Discussion

Skilled movement execution relies on precise connectivity between SCPN and their supraspinal as well as spinal targets. In this report, we identify that (1) axon targeting specificity by CBN is established from the initial stages of white matter axon extension, suggesting molecular mechanisms that limit CBN axon extension to supraspinal levels; and (2) molecular delineation of CBN vs. CSN in early development indicates a specification of these anatomically distinct projections. We further identify that the molecular differences between CBN and CSN can prospectively delineate these subpopulations and that these genes also exhibit temporal dynamics that correlate with the developmental time points of axon extension specificity at the respective levels of the neuraxis. Within lateral cortex, CBN are specifically identified by *Npy* expression, while cervical projecting CSN in lateral cortex are identified by *Cartpt* expression. This molecular delineation now provides, for the first time, molecular access to these subpopulations from early development into maturity. Our report is the first to identify a potential molecular program that may limit CBN axon extension to supraspinal levels.

### Cortico-brainstem axons are limited to supraspinal levels from initial phase of axon extension

Seminal work in the field had previously identified that SCPN throughout the neocortex extend exuberant axonal projections to the spinal cord early in development and that the specificity of axonal projections is then established via selective axonal pruning (Low et al., 2008; Luo & O’Leary, 2005; Martin, 2005). Our results now add an additional complexity to this model of circuit establishment. Our retrograde tracing analyses were performed at the earliest time points of initial axon extension to either the brainstem or the spinal cord. We establish that cortico-brainstem axons indeed are limited to supraspinal levels, and do not extend into the spinal cord from this earliest time of spinal axon extension in development. This indicates that there is already molecular specification that likely controls this initial axon extension specificity. Therefore, well before axonal pruning, axon extension establishes an earlier level of specificity in establishing segmental target diversity. Together with our prior work (Sahni, Shnider, et al., 2021), this suggests an early developmental molecular control over axon extension specificity in the white matter between distinct SCPN subpopulations.

### Molecular diversification of SCPN across multiple axes during development

Recently, there has been tremendous progress in identifying diversity of cell types in the developing and adult cerebral cortex using single-cell profiling (Bakken et al., 2016, 2021; La Manno et al., 2021; Loo et al., 2019; Rosenberg et al., 2018; Tasic, 2018; Tasic et al., 2016; Zeisel et al., 2015), including the adult primary motor cortex (Yao et al., 2021). Since epigenetic mechanisms can also mediate postmitotic acquisition of subtype identity (Harb et al., 2016), investigations have integrated gene expression analysis with chromatin accessibility via single-cell ATAC-seq (Armand et al., 2021; Preissl et al., 2018). Therefore, integrated cell-type atlases of the developing cerebral cortex are being established that also delineate developmental trajectories of distinct neocortical neuron subtypes (Di Bella et al., 2021; Heavner et al., 2020). While these are important studies that define the overall cellular diversity within the cerebral cortex, minority populations of neurons are often underrepresented in these datasets. CSN, for instance, constitute < 0.1 % of total cortical neurons (cortical neurons in mice = ∼14 million (Herculano-Houzel, 2009; Herculano-Houzel et al., 2006), CSN = ∼5000 (Bareyre et al., 2005)). Thus, datasets analyzing broad cortical diversity are limited in their ability to identify projection targeting and additional layers of diversity within SCPN. Further, developmental genes that control early specification are often absent in adult cortex.

Our prior work identified that SCPN in medial versus lateral cortex exhibit segmentally distinct projections, with lateral SCPN projecting to bulbar-cervical levels specifically (Sahni, Shnider, et al., 2021). Interestingly, in our single-cell RNA-seq data, cortical location provides one axis of SCPN molecular diversity with medial vs. lateral SCPN segregating into distinct clusters. Additionally, projection targeting diversity also contributes to SCPN molecular diversity, where CBN cluster together and CSN cluster together. However, the axis of projection targeting diversity is distinct from the axis that segregates SCPN along the cortical location. Together, these axes differentiate SCPN into medial vs. lateral residing CBN / CSN, respectively, with likely additional layers of diversity within each of these broad clusters. This further suggests that cortical location alone does not pre-specify axon extension specificity, which is consistent with our previous findings that bulbar-cervical and thoraco-lumbar projecting neurons can reside intermingled in the same cortical location (Sahni, Shnider, et al., 2021).

In our anatomical distinction between medial vs. lateral cortex, the primary somatosensory cortex (S1) is included in lateral cortex, while primary motor cortex (M1), as well as the transition zone between agranular motor cortex and granular sensory cortex is included in medial cortex (Sahni, Shnider, et al., 2021; Tennant et al., 2011). Our previous work identified that SCPN residing in medial cortex, which includes eventual M1 in the adult, consist of multiple distinct subpopulations: from cortico-brainstem, to corticospinal neurons projecting to cervical, thoracic, or lumbar spinal cord (Sahni, Shnider, et al., 2021). In our single-cell RNA-seq data, we find multiple, distinct clusters within medial cortex both for CBN and CSN, confirming previous reports as well as our anatomical data. Indeed, some of the clusters that were *post hoc* annotated as medial and spinal projecting showed a high expression of Crim1 and St6galnac5, which were previously shown to be specific for thoraco-lumbar projecting neurons (Sahni, Shnider, et al., 2021). The additionally identified spinal medial clusters might represent cervical-projecting CSN in medial cortex, however, top differentially expressed genes for these clusters have yet to be validated. In total, our dataset confirms that SCPN in medial cortex are more heterogenous and contain CBN, as well as distinct CSN subpopulations (cervical-, thoracic- and lumbar-projecting).

Using retrograde labeling, we find SCPN in lateral cortex are predominantly CBN, with a smaller subset of CSN, which are likely to be cervical-projecting (Sahni, Shnider, et al., 2021). The *post hoc* annotated spinal lateral cluster in our single-cell RNA-seq data was specifically defined by expression of *Cartpt*, which was previously identified to be expressed by bulbar-cervical projecting CSN in lateral cortex (Sahni, Shnider, et al., 2021). We now further identify and delineate that *Cartpt* is specific to cervical-projecting CSN, as it was excluded from CBN in the correlation analyses (Fig. 3G). Supporting this, *Cartpt* expression increases from P1 to P3, as corticospinal axons have crossed the brainstem-spinal transition zone to extend into the spinal cord. Recent work has identified that cervical-projecting CSN in lateral cortex can non-cell-autonomously regulate axon collateralization in the cervical gray matter by CSN residing in medial cortex. This non-cell-autonomous suppression is mediated by the proteoglycan *Lumican* (*Lum*). *Lum* expression by these neurons increases at later developmental times (Itoh et al., 2021). In line with this, we do not find *Lum*^+^ SCPN in our single-cell RNA-seq dataset. Our single-cell RNA-seq data indicates that these cervical-projecting CSN in lateral cortex are likely *Cartpt*^+^ at earlier time points. It will be interesting to investigate whether *Cartpt*^+^ CSN begin to express *Lum* at later developmental times, and to also identify additional regulators that control later aspects of corticospinal connectivity and circuit-level refinement.

In contrast, the brainstem lateral cluster is specified by high differential expression of *Npy*, a neuropeptide usually known to be expressed by cortical interneurons. *Npy* was recently described at postnatal day P1 in layer 5/6 callosal projection neurons, as well as corticofugal neurons, which encompass SCPN among others (Di Bella et al., 2021). Using expression analyses, as well as conditional labeling in Npy-IRES-Cre knock-in reporter mice, we establish that *Npy* is specifically expressed by CBN in lateral cortex. This highly restricted *Npy* expression suggests an additional level of post-mitotic differentiation of SCPN into CBN vs. CSN that occurs early during SCPN axon extension. Further, *Npy* expression levels decline from P1 to P3, which is the time point at which CBN limit their axon extension to supraspinal levels and do not extend axons beyond the brainstem-spinal transition zone into the spinal cord. These temporal changes likely reflect the dynamic molecular regulation of this initial axon extension specificity. Our dataset provides additional candidate genes that delineate CBN from CSN in early postnatal development and potentially identify novel molecular controls over this developmental axon extension specificity.

### Therapeutic potential of cortico-brainstem-spinal pathways in neurological disorders

The brainstem has been widely recognized as a key integration and processing hub between “upper” motor centers and spinal circuits involved in execution of movements (Arber & Costa, 2018; Lemon, 2008; Ruder & Arber, 2019). Cortical input into the brainstem may fine-tune movement execution and support motor planning (Economo et al., 2018; Svoboda & Li, 2018). After lesions to the motor cortex, such as in cortical stroke, the brainstem may provide an additional relay station for compensatory recovery. Indeed, it has been shown that both brainstem-spinal as well as cortico-brainstem innervation increased after a large cortical stroke both spontaneously and by the use of rehabilitative training (Bachmann et al., 2014; Ishida et al., 2016), and that a second cortical lesion that disrupts these new connections is detrimental to the recovered motor behavior. Understanding how the specificity of this cortico-brainstem-spinal innervation is established early in development might also help inform how this circuit is remodeled in instances of disease or injury.

Taken together, this work establishes that there are multiple choice points along the rostro-caudal extent of the neuraxis – the brainstem-spinal transition zone, the cervical- thoracic transition zone – that establish the initial segmental targeting specificity of subcerebral connectivity. In this study, we address early molecular delineation of SCPN axon extension specificity into the brainstem vs. the spinal cord at initial time points of axon extension. We identify molecular delineation of SCPN projection targeting diversity within the lateral cortex, that molecularly distinguishes CBN (*Npy*^+^) from cervical projecting CSN (*Cartpt*^+^).

This early molecular differentiation over SCPN axon extension specificity is likely one of several developmental processes to establish distinct SCPN subsets that will eventually participate in functionally distinct and relevant circuits. Future investigations can leverage these molecular differences, e.g., Npy-IRES-Cre transgenic mice, to analyze these molecularly defined SCPN subpopulations from early development into maturity at a molecular, circuit, and functional level. Our work therefore provides the foundation to identify novel molecular mechanisms, including activity-dependent processes that control subsequent aspects of their distinct differentiation and connectivity. Further, these molecular tools will enable functional investigations into the contributions of these defined subsets to distinct aspects of skilled motor function. Together, our findings identifying this early molecular diversity of SCPN will provide useful insights into the development, plasticity, and function of this critical motor control circuit.

## Methods

### Mice

Wild-type CD1 mice were obtained from Charles River Laboratories (Wilmington, MA). The day of birth was designated at P0. B6.Cg-Npy^tm1(cre)Zman^/J (herein called Npy-IRES-Cre) mice were obtained from Jackson Laboratories (Stock No.: 027851) and have an IRES and a Cre recombinase cassette inserted into the 3’ UTR of the Npy locus downstream of the stop codon. All mouse studies were performed in accordance with institutional and federal guidelines and were approved by the Weill Cornell Medical College Institutional animal care and use committee.

### Retrograde labeling of layer V neurons

In all experiments, SCPN (CBN + CSN) or CSN only were retrogradely labeled by bilateral injection of Cholera Toxin B subunit (CTB; Thermo Scientific) into either cerebral peduncles (at 3 injection sites, 161nl injected at each site in 23nl increments; total of 483 nl) or into the spinal segment C1/C2 (both sides of the midline, total of 207 nl), respectively, using ultrasound-guided backscatter microscopy (Vevo 2100; VisualSonics, Toronto, Canada) via a pulled glass micropipette with a nanojector (Nanoject II, Drummond Scientific, Broomall, PA) at a rate of 23 nl/ second. Injections were performed at the time point of initial axon extension (P0 at cerebral peduncles, P2 at cervical spinal cord) unless stated otherwise. For AAV-mediated retrograde labeling in the Npy-IRES-Cre mice rAAV-CAG-Flex-eGFP-WPRE and rAAV-CAG-tdTomato were co-injected. The rAAV-pCAG-FLEX-EGFP-WPRE was a gift from Hongkui Zeng (Addgene plasmid #51502-AAVrg; RRID:Addgene_51502 (Oh et al., 2014)) and the rAAV-CAG-tdTomato (codon diversified) was a gift from Edward Boyden (Addgene plasmid #59462-AAVrg; RRID:Addgene_59462).

### Quantification of retrogradely labeled cells

Following retrograde labeling of SCPN or CSN with CTB, mice were anesthetized on ice and transcardially perfused with 4% paraformaldehyde (PFA) at P4. Brains were dissected, post-fixed in 4% PFA overnight and stored at 4°C in PBS until further use. Tissue was cryopreserved in 30% sucrose overnight and frozen in Tissue-Tek OCT Compound (Sakura Finetek, Torrance, CA). Coronal brain sections (50 µm) were imaged in 4 µm z-stacks at 20x on a Zeiss Axioimager M2 microscope using the Stereo Investigator software (MBF Biosciences). A single plane image from the z-stacks was obtained using the Deep Focus tool in the Neurolucida software (MBF Biosciences). Retrogradely labeled neurons were counted manually in Neurolucida at 4 different levels (1.95mm, 2.67mm, 3.27mm, 4.23mm to Bregma). Blinding of the observers was not possible due to obvious differences in the groups. For each coronal section, 5 bins were placed over the width of the hemisphere to determine the cingulate (1 bin) vs medial (2 bins) vs lateral (2 bins) cortex distinction.

Following AAV-mediated retrograde conditional labeling, Npy-IRES-Cre mice were perfused at P14 as described above. Coronal brain sections (50 µm) were imaged at 10x on a Zeiss Axioimager M2 microscope and labeled neurons (Npy^+^, eGFP^+^) were detected using AMaSiNe (Song et al., 2020) following the recommended protocol (vsnnlab, 2020/2021) with minor manual edits to obtain a 3D representation of the location of traced neurons.

### *In Situ* Hybridization/RNAScope

After retrograde labeling of SCPN or CSN, CD1 mice were transcardially perfused with cold 4% PFA at P3. Brains were dissected and post-fixed overnight. After cryopreservation in 30% sucrose overnight, the tissues were directly embedded in Tissue-Tek OCT Compound prior to sectioning. Coronal sections (30µm) were collected on slide and processed using the RNAScope Multiplex Fluorescent v2 kit (Advanced Cell Diagnostics, 323100) for gene expression of Cartpt (Advanced Cell Diagnostics, 432001-C2) or Npy (ACD Bio, 313321-C3). The CTB signal was detected by combining the RNAScope Multiplex Fluorescent v2 assay with immunofluorescence using a rabbit anti-CTB primary antibody (Abcam, ab34992, 1:200). The integrated RNA protein co-detection workflow was followed using the RNA-Protein Co-detection Ancillary Kit (Advanced Cell Diagnostics, 323180) as per manufacturer’s instructions.

### Fluorescent activated cell sorting (FACS)

To sort labeled SCPN, tissue of retrogradely labeled mice was collected at P1 (SCPN group only) or P3 (SCPN or CSN group). All procedures were done rapidly and using cold solutions to minimize time between euthanasia and cell collection. Briefly, brains were dissected and sliced into 800 µm thick coronal sections which were subsequently processed under a fluorescent dissection scope (Nikon SMZ18) to extract cortex containing labeled cells. A single cell suspension of the samples was obtained by enzymatic (15min, 37°C) and mechanical dissociation (gentle trituration using rounded glass Pasteur pipettes of decreasing diameter (600um, 300um)) followed by filtration through a 40µm mesh (Biologix, 15-1040-1).

SCPN were FACS-purified from this single cell suspension using a WOLF flow sorter (Nanocellect Biomedical) with standard settings (threshold for cell size adapted to 30.000). To decrease contamination and increase sorting efficacy, cells were initially sorted in a high concentration of ∼6 × 10^6^ cells/ml followed by a second sort of the first round of sorted cells (lower concentration, ∼2 × 10^5^ cells/ml). This allowed for enrichment of CTB 555 labeled cells in a timely manner. We collected ∼15.000 cells in 5ml per sort. Cells were enriched for downstream processing by centrifugation for 10min at 80g and processed directly for single-cell RNA-Seq.

### Single-cell RNA sequencing

Each sample (P1.SCPN, P3.SCPN and P3.CSN) was sequenced using the Chromium 10x Single Cell 3’ pipeline following the standard protocol. For the SCPN samples, experiments were repeated on two separate occasions (biological replicates) and two samples were collected each time (technical replicate). For the CSN samples, two technical replicates were collected. Briefly, single cell suspension of 5000 to 10000 cells were loaded onto a Chromium Chip B / G and processed using the standard protocol of the Chromium Next GEM Single Cell 3’ GEM and Gel bead Kits v3 and v3.1, respectively. Libraries were prepared using the Chromium Next GEM Single Cell 3’ Library Kit v3.1 and the Chromium i7 Multiplex barcodes. The sequencing libraries (n=11, P3.CSN (2), P3.SCPN (5), P1.SCPN (4)) were evaluated for quality on the Agilent TapeStation (Agilent Technologies, Palo Alto, CA, USA), quantified using Qubit 2.0 Fluorometer (Invitrogen, Carlsbad, CA) and pooled libraries were quantified using qPCR (Applied Biosystems, Carlsbad, CA, USA). The pooled sequencing libraries were clustered on 5 lanes of a flow cell and loaded on the Illumina HiSeq instrument (4000 or equivalent) according to manufacturer’s instructions. The samples were sequenced in a configuration compatible with the recommended guidelines as outlined by 10X Genomics (2×150bp configuration, with 8 bp single index).

### Bioinformatics

Single-cell RNA-seq data were processed as recommended by Amezquite et al. (2020). In brief, raw read files were processed and aligned to the mouse reference, *mm10* (GENCODE vM23/Ensembl 98) using 10x Genomics Cell Ranger 6.0.1 *count* and *aggr* (Zheng et al., 2017). Using established R packages and custom-written code, cells with low read counts (empty droplets) or high numbers of mitochondrial gene products (±3x than median absolute deviation) were removed. Size-factor normalized logcounts were obtained (A. T. L. Lun, Bach, et al., 2016; McCarthy et al., 2017) and batch-corrected (Haghverdi et al., 2018). After integration of three sample types (P1 SCPN, P3 SCPN, P3 CSN) using the top 2000 most variable genes with min. mean normalized expression of 0.001 (A. Lun, 2019), dimensionality reduction (UMAP) was done on the batch-corrected log counts. The dataset was annotated with singleR (Aran et al., 2019; A. T. L. Lun et al., 2020) using previously published single-cell data of the P0 mouse cortex to subset our dataset to Layer V neurons (Loo et al., 2019). Following *in silico* extraction of layer V neurons, projection targeting specificity was defined as follows: clusters that were only present in the SCPN but not the CSN samples were annotated as “brainstem projecting”, while overlapping clusters were annotated as “spinal projecting”. Medial vs. lateral cortical location was annotated with singleR based on microarray data obtained in a previous study (Sahni, Shnider, et al., 2021) for the P1 sample (matching the time point of the microarray). The dataset was subsequently clustered with graph-based clustering (k = 50 (A. T. L. Lun, McCarthy, et al., 2016)), resulting clusters were manually annotated based on projection targeting and cortical location and marker genes between clusters were obtained using *findMarker* function in Seurat v4.0 (Hao et al., 2021).

### Statistical analysis

Statistical analysis other than those related to the single-cell RNA-seq data was performed with Prism 7.0 (GraphPad Software). For the quantification of retrogradely labeled neurons at time points of initial axon extension, mixed-effect models were used within each bin followed by Šídák’s multiple comparisons test. No statistical methods were used to pre-determine sample sizes. In the bar graph, dots represent individual animals. The threshold for significance for all experiments was set at \**p* < 0.05. Smaller *p* values were represented as \*\**p* < 0.01 and \*\*\**p* < 0.001. In bar graphs, all data are plotted as mean ± SEM. In box plot graphs, data are represented as median ± 25th percentile (box) and min/max (whiskers), and individual dots represent single cells. Comparison between groups was done using students t-test.

## Acknowledgements

We thank members of the Hollis and Sahni Labs for useful discussions and comments on the manuscript. We thank Michał Bonar from NanoCellect for providing excellent technical support and protocol advice for FACS purification. This research was supported by NIH (NTRAIN/NICHD K12HD093427), and grants from the Wings for Life - Spinal Cord Injury Foundation and the Craig Neilsen Foundation, as well as additional infrastructure support from the Burke Foundation to V.S.. J.K. was partially supported by a postdoctoral fellowship from the Swiss National Science Foundation (SNSF, P2EZP3_191858).

## Supplementary Figures

**Supplemental Fig 1.**
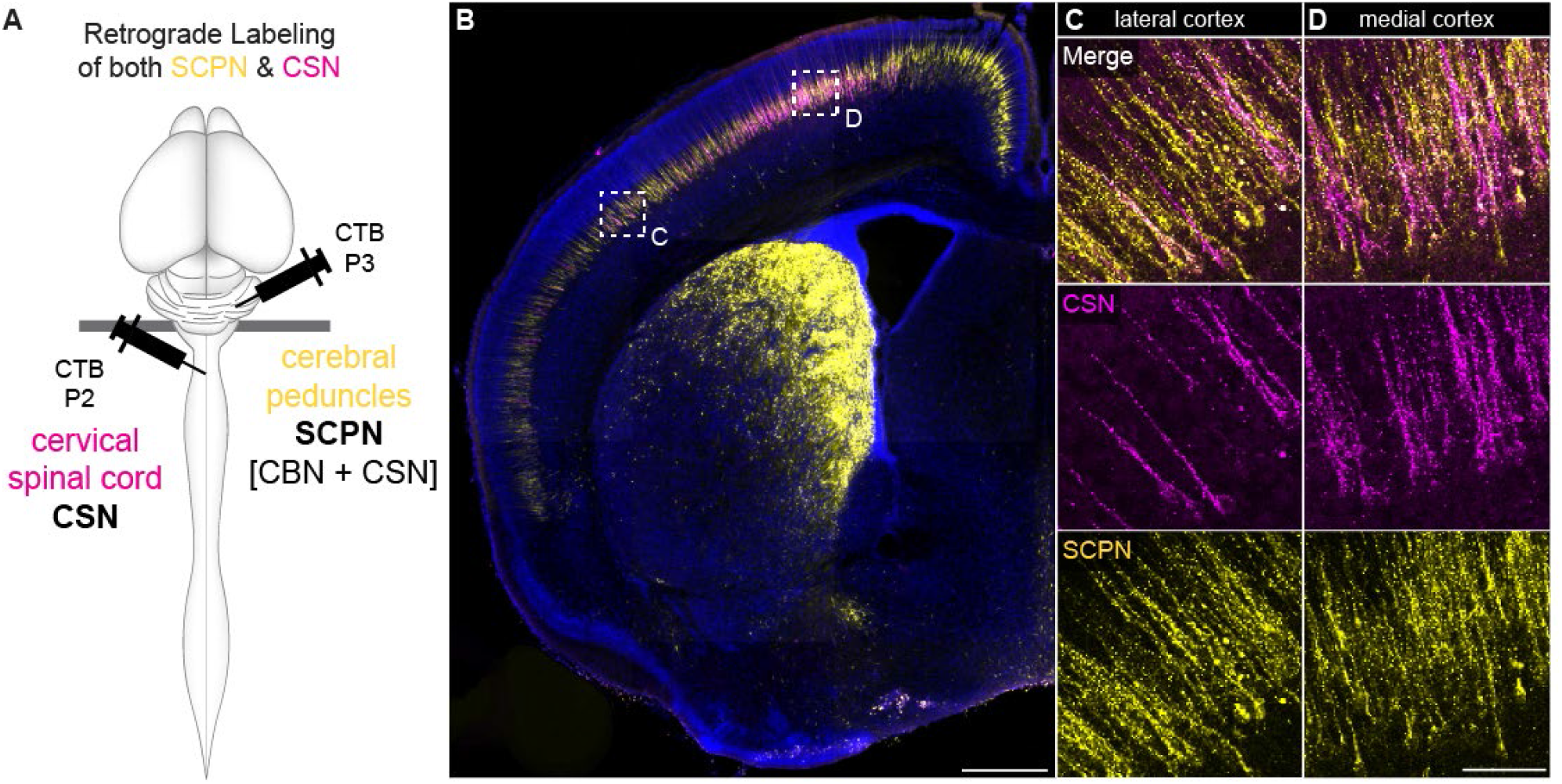
Establishment of the brainstem-spinal transition zone in early development. (A) Experimental scheme, CSN or SCPN (CBN + CSN) were retrogradely labeled by injection into the spinal cord (CSN) or at the level of the cerebral peduncles (SCPN), respectively, in the same animal. For this, we first injected into the spinal cord at P2, followed by injection of a second, distinct tracer into the cerebral peduncles at P3 to minimize injection artifacts in labeling. (B) Coronal section of a retrogradely labeled brain at P4 showing all SCPN (yellow), and CSN, as a subset of all SCPN (magenta). Labeled SCPN can be seen across both medial and lateral cortex in layer V. Scale bar, 500µm. (C) In lateral cortex, few of the labeled SCPN colocalize with the CSN label; thus, the majority of labeled cells are CBN. (D) In medial cortex, the number of SCPN that are co-labeled by the spinal tracer (CSN) is higher, the population of SCPN is more mixed. Scale bar, 50µm.

**Supplemental Fig 2.**
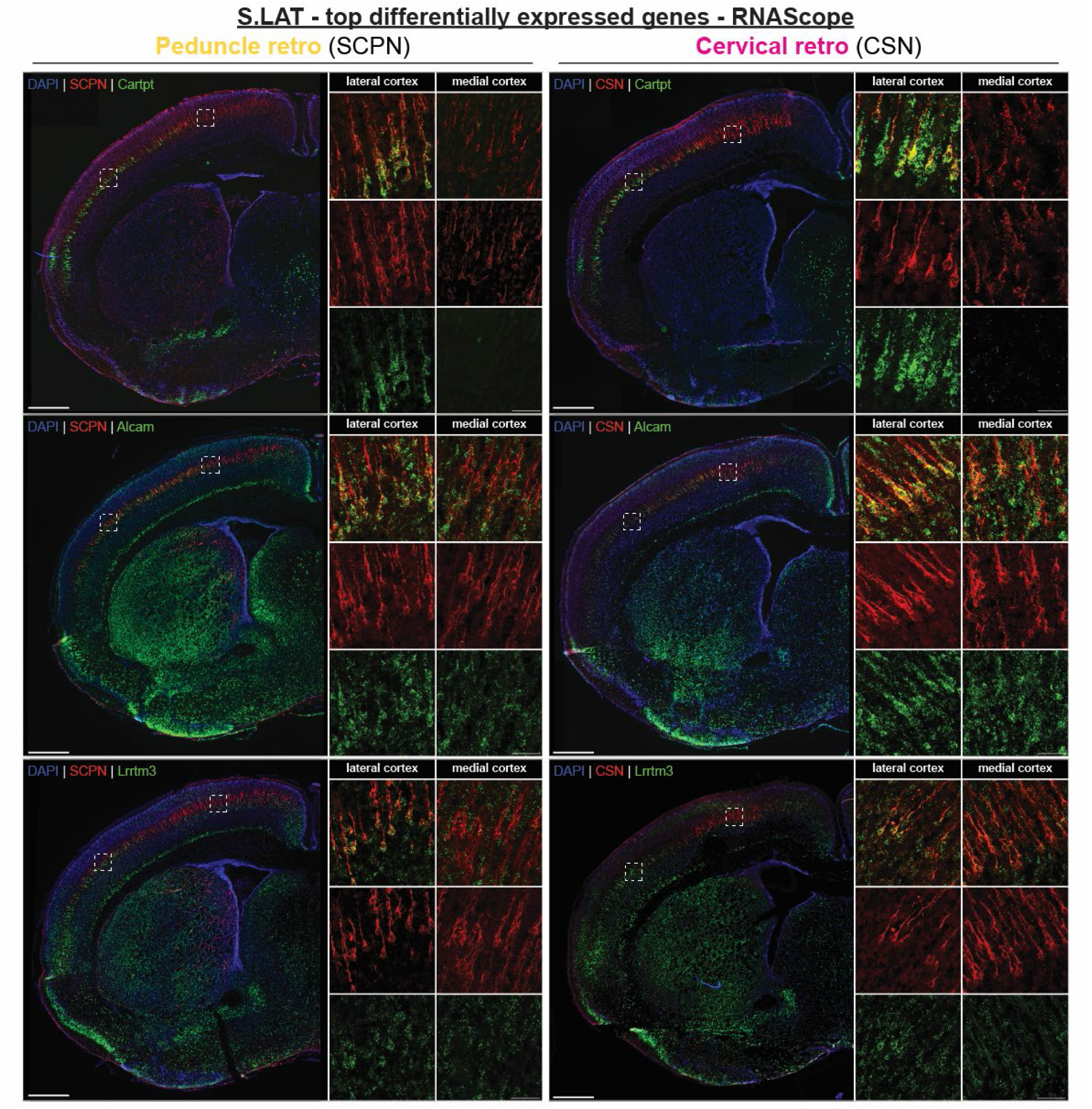
Cartpt, Alcam and Lrrtm3 are expressed in lateral cortex by corticospinal neurons. In-situ hybridization for Cartpt (top row), Alcam (middle row) and Lrrtm3 (bottom row). Co-localization with retrogradely labeled SCPN (left) and CSN (right) shows specificity of all genes to the lateral cortex. Co-localization of RNAScope signal with retrogradely labeled CSN suggests that these genes are expressed in cervical projecting CSN in lateral cortex at P3.

**Supplemental Fig 3.**
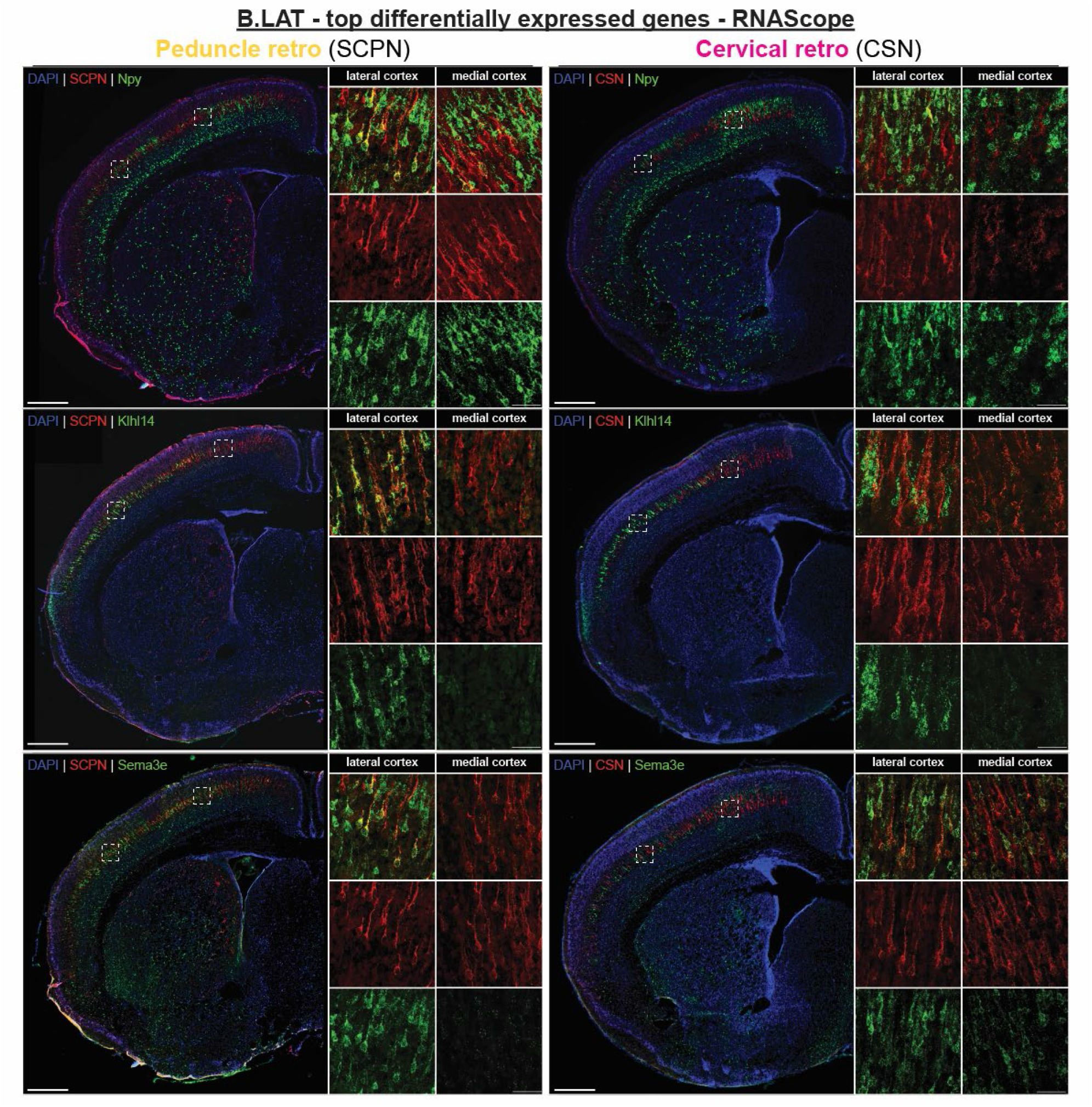
Npy, Klhl14 and Sema3e are expressed specifically in CBN and not CSN in lateral cortex. In-situ hybridization for Npy (top row), Klhl14 (middle row) and Sema3e (bottom row). Co-localization with retrogradely labeled SCPN (left) and not CSN (right) shows that these genes are expressed by CBN at P3.

## References

Amezquita, R. A., Lun, A. T. L., Becht, E., Carey, V. J., Carpp, L. N., Geistlinger, L., Marini, F., Rue-Albrecht, K., Risso, D., Soneson, C., Waldron, L., Pagès, H., Smith, M. L., Huber, W., Morgan, M., Gottardo, R., & Hicks, S. C. (2020). Orchestrating single-cell analysis with Bioconductor. Nature Methods, 17(2), 137–145. https://doi.org/10.1038/s41592-019-0654-x

Aran, D., Looney, A. P., Liu, L., Wu, E., Fong, V., Hsu, A., Chak, S., Naikawadi, R. P., Wolters, P. J., Abate, A. R., Butte, A. J., & Bhattacharya, M. (2019). Reference-based analysis of lung single-cell sequencing reveals a transitional profibrotic macrophage. Nature Immunology, 20(2), 163–172. https://doi.org/10.1038/s41590-018-0276-y

Arber, S., & Costa, R. M. (2018). Connecting neuronal circuits for movement. Science, 360(6396), 1403–1404. https://doi.org/10.1126/science.aat5994

Arlotta, P., Molyneaux, B. J., Chen, J., Inoue, J., Kominami, R., & MacKlis, J. D. (2005). Neuronal subtype-specific genes that control corticospinal motor neuron development in vivo. Neuron, 45(2), 207–221. https://doi.org/10.1016/j.neuron.2004.12.036

Armand, E. J., Li, J., Xie, F., Luo, C., & Mukamel, E. A. (2021). Single-Cell Sequencing of Brain Cell Transcriptomes and Epigenomes. Neuron, 109(1), 11–26. https://doi.org/10.1016/j.neuron.2020.12.010

Bachmann, L. C., Lindau, N. T., Felder, P., & Schwab, M. E. (2014). Sprouting of Brainstem-Spinal Tracts in Response to Unilateral Motor Cortex Stroke in Mice. Journal of Neuroscience, 34(9), 3378–3389. https://doi.org/10.1523/jneurosci.4384-13.2014

Bakken, T. E., Jorstad, N. L., Hu, Q., Lake, B. B., Tian, W., Kalmbach, B. E., Crow, M., Hodge, R. D., Krienen, F. M., Sorensen, S. A., Eggermont, J., Yao, Z., Aevermann, B. D., Aldridge, A. I., Bartlett, A., Bertagnolli, D., Casper, T., Castanon, R. G., Crichton, K., … Lein, E. S. (2021). Comparative cellular analysis of motor cortex in human, marmoset and mouse. Nature, 598(7879), 111–119. https://doi.org/10.1038/s41586-021-03465-8

Bakken, T. E., Miller, J. A., Ding, S.-L., Sunkin, S. M., Smith, K. A., Ng, L., Szafer, A., Dalley, R. A., Royall, J. J., Lemon, T., Shapouri, S., Aiona, K., Arnold, J., Bennett, J. L., Bertagnolli, D., Bickley, K., Boe, A., Brouner, K., Butler, S., … Lein, E. S. (2016). A comprehensive transcriptional map of primate brain development. Nature, 535(7612), 367–375. https://doi.org/10.1038/nature18637

Bareyre, F. M., Kerschensteiner, M., Misgeld, T., & Sanes, J. R. (2005). Transgenic labeling of the corticospinal tract for monitoring axonal responses to spinal cord injury. Nature Medicine, 11(12), 1355–1360. https://doi.org/10.1038/nm1331

Cederquist, G. Y., Azim, E., Shnider, S. J., Padmanabhan, H., & Macklis, J. D. (2013). Lmo4 Establishes Rostral Motor Cortex Projection Neuron Subtype Diversity. Journal of Neuroscience, 33(15), 6321–6332. https://doi.org/10.1523/jneurosci.5140-12.2013

Chen, B., Schaevitz, L. R., & McConnell, S. K. (2005). Fezl regulates the differentiation and axon targeting of layer 5 subcortical projection neurons in cerebral cortex. Proceedings of the National Academy of Sciences, 102(47), 17184–17189. https://doi.org/10.1073/pnas.0508732102

Chen, J. G., Rašin, M. R., Kwan, K. Y., & Šestan, N. (2005). Zfp312 is required for subcortical axonal projections and dendritic morphology of deep-layer pyramidal neurons of the cerebral cortex. Proceedings of the National Academy of Sciences of the United States of America, 102(49), 17792–17797. https://doi.org/10.1073/pnas.0509032102

Conner, J. M., Bohannon, A., Igarashi, M., Taniguchi, J., Baltar, N., & Azim, E. (2021). Modulation of tactile feedback for the execution of dexterous movement. Science, 374(6565), 316–323. https://doi.org/10.1126/science.abh1123

Di Bella, D. J., Habibi, E., Stickels, R. R., Scalia, G., Brown, J., Yadollahpour, P., Yang, S. M., Abbate, C., Biancalani, T., Macosko, E. Z., Chen, F., Regev, A., & Arlotta, P. (2021). Molecular logic of cellular diversification in the mouse cerebral cortex. Nature, 595(7868), 554–559. https://doi.org/10.1038/s41586-021-03670-5

Economo, M. N., Clack, N. G., Lavis, L. D., Gerfen, C. R., Svoboda, K., Myers, E. W., & Chandrashekar, J. (2016). A platform for brain-wide imaging and reconstruction of individual neurons. ELife, 5, e10566. https://doi.org/10.7554/eLife.10566

Economo, M. N., Viswanathan, S., Tasic, B., Bas, E., Winnubst, J., Menon, V., Graybuck, L. T., Nguyen, T. N., Smith, K. A., Yao, Z., Wang, L., Gerfen, C. R., Chandrashekar, J., Zeng, H., Looger, L. L., & Svoboda, K. (2018). Distinct descending motor cortex pathways and their roles in movement. Nature, 563(7729), 79–84. https://doi.org/10.1038/s41586-018-0642-9

Esposito, M. S., Capelli, P., & Arber, S. (2014). Brainstem nucleus MdV mediates skilled forelimb motor tasks. Nature, 508(7496), 351–356. https://doi.org/10.1038/nature13023

Fame, R. M., MacDonald, J. L., & Macklis, J. D. (2011). Development, specification, and diversity of callosal projection neurons. Trends in Neurosciences, 34(1), 41–50. https://doi.org/10.1016/j.tins.2010.10.002

Franco, S. J., & Müller, U. (2013). Shaping Our Minds: Stem and Progenitor Cell Diversity in the Mammalian Neocortex. Neuron, 77(1), 19–34. https://doi.org/10.1016/j.neuron.2012.12.022

Greig, L. C., Woodworth, M. B., Galazo, M. J., Padmanabhan, H., & Macklis, J. D. (2013). Molecular logic of neocortical projection neuron specification, development and diversity. Nature Reviews Neuroscience, 14(11), 755–769. https://doi.org/10.1038/nrn3586

Greig, L. C., Woodworth, M. B., Greppi, C., & Macklis, J. D. (2016). Ctip1 Controls Acquisition of Sensory Area Identity and Establishment of Sensory Input Fields in the Developing Neocortex. Neuron, 90(2), 261–277. https://doi.org/10.1016/j.neuron.2016.03.008

Haghverdi, L., Lun, A. T. L., Morgan, M. D., & Marioni, J. C. (2018). Batch effects in single-cell RNA-sequencing data are corrected by matching mutual nearest neighbors. Nature Biotechnology, 36(5), 421–427. https://doi.org/10.1038/nbt.4091

Han, W., Kwan, K. Y., Shim, S., Lam, M. M. S., Shin, Y., Xu, X., Zhu, Y., Li, M., & Šestan, N. (2011). TBR1 directly represses Fezf2 to control the laminar origin and development of the corticospinal tract. Proceedings of the National Academy of Sciences, 108(7), 3041–3046. https://doi.org/10.1073/pnas.1016723108

Hao, Y., Hao, S., Andersen-Nissen, E., Mauck, W. M., Zheng, S., Butler, A., Lee, M. J., Wilk, A. J., Darby, C., Zager, M., Hoffman, P., Stoeckius, M., Papalexi, E., Mimitou, E. P., Jain, J., Srivastava, A., Stuart, T., Fleming, L. M., Yeung, B., … Satija, R. (2021). Integrated analysis of multimodal single-cell data. Cell, 184(13), 3573-3587.e29. https://doi.org/10.1016/j.cell.2021.04.048

Harb, K., Magrinelli, E., Nicolas, C. S., Lukianets, N., Frangeul, L., Pietri, M., Sun, T., Sandoz, G., Grammont, F., Jabaudon, D., Studer, M., & Alfano, C. (2016). Area-specific development of distinct projection neuron subclasses is regulated by postnatal epigenetic modifications. ELife, 5, e09531. https://doi.org/10.7554/eLife.09531

Heavner, W. E., Ji, S., Notwell, J. H., Dyer, E. S., Tseng, A. M., Birgmeier, J., Yoo, B., Bejerano, G., & McConnell, S. K. (2020). Transcription factor expression defines subclasses of developing projection neurons highly similar to single-cell RNA-seq subtypes. Proceedings of the National Academy of Sciences, 117(40), 25074–25084. https://doi.org/10.1073/pnas.2008013117

Herculano-Houzel, S. (2009). The human brain in numbers: A linearly scaled-up primate brain. Frontiers in Human Neuroscience, 3. https://www.frontiersin.org/article/10.3389/neuro.09.031.2009

Herculano-Houzel, S., Mota, B., & Lent, R. (2006). Cellular scaling rules for rodent brains. Proceedings of the National Academy of Sciences, 103(32), 12138–12143. https://doi.org/10.1073/pnas.0604911103

Ishida, A., Isa, K., Umeda, T., Kobayashi, K., Kobayashi, K., Hida, H., & Isa, T. (2016). Causal Link between the Cortico-Rubral Pathway and Functional Recovery through Forced Impaired Limb Use in Rats with Stroke. The Journal of Neuroscience, 36(2), 455–467. https://doi.org/10.1523/JNEUROSCI.2399-15.2016

Itoh, Y., Sahni, V., Shnider, S., & Macklis, J. (2021). Lumican regulates cervical corticospinal axon collateralization via non-autonomous crosstalk between distinct corticospinal neuron subpopulations. BioRxiv, 617.

Joshi, P. S., Molyneaux, B. J., Feng, L., Xie, X., Macklis, J. D., & Gan, L. (2008). Bhlhb5 Regulates the Postmitotic Acquisition of Area Identities in Layers II-V of the Developing Neocortex. Neuron, 60(2), 258–272. https://doi.org/10.1016/j.neuron.2008.08.006

Kaiser, J., Maibach, M., Salpeter, I., Hagenbuch, N., Souza, V. B. C., Robinson, M. D., & Schwab, M. E. (2019). The spinal transcriptome after cortical stroke—In search of molecular factors regulating spontaneous recovery in the spinal cord. The Journal of Neuroscience, 39(24), 2571–18. https://doi.org/10.1523/JNEUROSCI.2571-18.2019

Karagiannis, A., Gallopin, T., David, C., Battaglia, D., Geoffroy, H., Rossier, J., Hillman, E. M. C., Staiger, J. F., & Cauli, B. (2009). Classification of NPY-Expressing Neocortical Interneurons. Journal of Neuroscience, 29(11), 3642–3659. https://doi.org/10.1523/JNEUROSCI.0058-09.2009

Kwan, K. Y., Lam, M. M. S., Krsnik, Ž., Kawasawa, Y. I., Lefebvre, V., & Šestan, N. (2008). SOX5 postmitotically regulates migration, postmigratory differentiation, and projections of subplate and deep-layer neocortical neurons. Proceedings of the National Academy of Sciences, 105(41), 16021–16026. https://doi.org/10.1073/pnas.0806791105

La Manno, G., Siletti, K., Furlan, A., Gyllborg, D., Vinsland, E., Mossi Albiach, A., Mattsson Langseth, C., Khven, I., Lederer, A. R., Dratva, L. M., Johnsson, A., Nilsson, M., Lönnerberg, P., & Linnarsson, S. (2021). Molecular architecture of the developing mouse brain. Nature, 596(7870), 92–96. https://doi.org/10.1038/s41586-021-03775-x

Lai, T., Jabaudon, D., Molyneaux, B. J., Azim, E., Arlotta, P., Menezes, J. R. L., & Macklis, J. D. (2008). SOX5 Controls the Sequential Generation of Distinct Corticofugal Neuron Subtypes. Neuron, 57(2), 232–247. https://doi.org/10.1016/j.neuron.2007.12.023

Lemon, R. N. (2008). Descending Pathways in Motor Control. Annual Review of Neuroscience, 31(1), 195–218. https://doi.org/10.1146/annurev.neuro.31.060407.125547

Leone, D. P., Heavner, W. E., Ferenczi, E. A., Dobreva, G., Huguenard, J. R., Grosschedl, R., & McConnell, S. K. (2015). Satb2 Regulates the Differentiation of Both Callosal and Subcerebral Projection Neurons in the Developing Cerebral Cortex. Cerebral Cortex, 25(10), 3406–3419. https://doi.org/10.1093/cercor/bhu156

Leone, D. P., Srinivasan, K., Chen, B., Alcamo, E., & McConnell, S. K. (2008). The determination of projection neuron identity in the developing cerebral cortex. Current Opinion in Neurobiology, 18(1), 28–35. https://doi.org/10.1016/j.conb.2008.05.006

Levine, A. J., Lewallen, K. A., & Pfaff, S. L. (2012). Spatial organization of cortical and spinal neurons controlling motor behavior. Current Opinion in Neurobiology, 5. https://doi.org/10.1016/j.conb.2012.07.002

Lodato, S., Molyneaux, B. J., Zuccaro, E., Goff, L. A., Chen, H. H., Yuan, W., Meleski, A., Takahashi, E., Mahony, S., Rinn, J. L., Gifford, D. K., & Arlotta, P. (2014). Gene co-regulation by Fezf2 selects neurotransmitter identity and connectivity of corticospinal neurons. Nature Neuroscience, 17(8), 1046–1054. https://doi.org/10.1038/nn.3757

Lodato, S., Shetty, A. S., & Arlotta, P. (2015). Cerebral cortex assembly: Generating and reprogramming projection neuron diversity. Trends in Neurosciences, 38(2), 117–125. https://doi.org/10.1016/j.tins.2014.11.003

Loo, L., Simon, J. M., Xing, L., McCoy, E. S., Niehaus, J. K., Guo, J., Anton, E. S., & Zylka, M. J. (2019). Single-cell transcriptomic analysis of mouse neocortical development. Nature Communications, 10(1), 134. https://doi.org/10.1038/s41467-018-08079-9

Low, L. K., Liu, X.-B., Faulkner, R. L., Coble, J., & Cheng, H.-J. (2008). Plexin signaling selectively regulates the stereotyped pruning of corticospinal axons from visual cortex. Proceedings of the National Academy of Sciences, 105(23), 8136–8141. https://doi.org/10.1073/pnas.0803849105

Lun, A. (2019, December 24). A description of the theory behind the fastMNN algorithm. https://marionilab.github.io/FurtherMNN2018/theory/description.html

Lun, A. T. L., Andrews, J. M., Dundar, F., & Bunis, D. (2020, June 14). Using SingleR to annotate single-cell RNA-seq data. https://www.bioconductor.org/packages/release/bioc/vignettes/SingleR/inst/doc/SingleR.html

Lun, A. T. L., Bach, K., & Marioni, J. C. (2016). Pooling across cells to normalize single-cell RNA sequencing data with many zero counts. Genome Biology, 17(1), 75. https://doi.org/10.1186/s13059-016-0947-7

Lun, A. T. L., McCarthy, D. J., & Marioni, J. C. (2016). A step-by-step workflow for low-level analysis of single-cell RNA-seq data with Bioconductor. F1000Research, 5, 2122. https://doi.org/10.12688/f1000research.9501.2

Luo, L., & O’Leary, D. D. M. (2005). Axon Retraction and Degeneration in Development and Disease. Annual Review of Neuroscience, 28(1), 127–156. https://doi.org/10.1146/annurev.neuro.28.061604.135632

Martin, J. H. (2005). The corticospinal system: From development to motor control. Neuroscientist, 11(2), 161–173. https://doi.org/10.1177/1073858404270843

McCarthy, D. J., Campbell, K. R., Lun, A. T. L., & Wills, Q. F. (2017). Scater: Pre-processing, quality control, normalization and visualization of single-cell RNA-seq data in R. Bioinformatics, 33(8), 1179–1186. https://doi.org/10.1093/bioinformatics/btw777

McKenna, W. L., Betancourt, J., Larkin, K. A., Abrams, B., Guo, C., Rubenstein, J. L. R., & Chen, B. (2011). Tbr1 and Fezf2 Regulate Alternate Corticofugal Neuronal Identities during Neocortical Development. Journal of Neuroscience, 31(2), 549–564. https://doi.org/10.1523/JNEUROSCI.4131-10.2011

McKenna, W. L., Ortiz-Londono, C. F., Mathew, T. K., Hoang, K., Katzman, S., & Chen, B. (2015). Mutual regulation between Satb2 and Fezf2 promotes subcerebral projection neuron identity in the developing cerebral cortex. Proceedings of the National Academy of Sciences, 112(37), 11702–11707. https://doi.org/10.1073/pnas.1504144112

Milstein, A. D., Bloss, E. B., Apostolides, P. F., Vaidya, S. P., Dilly, G. A., Zemelman, B. V., & Magee, J. C. (2015). Inhibitory Gating of Input Comparison in the CA1 Microcircuit. Neuron, 87(6), 1274–1289. https://doi.org/10.1016/j.neuron.2015.08.025

Molyneaux, B. J., Arlotta, P., Hirata, T., Hibi, M., & Macklis, J. D. (2005). Fezl is required for the birth and specification of corticospinal motor neurons. Neuron, 47(6), 817–831. https://doi.org/10.1016/j.neuron.2005.08.030

Molyneaux, B. J., Arlotta, P., Menezes, J. R. L., & Macklis, J. D. (2007). Neuronal subtype specification in the cerebral cortex. Nature Reviews Neuroscience, 8(6), 427–437. https://doi.org/10.1038/nrn2151

Moreno-Lopez, Y., Bichara, C., Delbecq, G., Isope, P., & Cordero-Erausquin, M. (2021). The corticospinal tract primarily modulates sensory inputs in the mouse lumbar cord. ELife, 10, e65304. https://doi.org/10.7554/eLife.65304

Oh, S. W., Harris, J. A., Ng, L., Winslow, B., Cain, N., Mihalas, S., Wang, Q., Lau, C., Kuan, L., Henry, A. M., Mortrud, M. T., Ouellette, B., Nguyen, T. N., Sorensen, S. A., Slaughterbeck, C. R., Wakeman, W., Li, Y., Feng, D., Ho, A., … Zeng, H. (2014). A mesoscale connectome of the mouse brain. Nature, 508(7495), 207–214. https://doi.org/10.1038/nature13186

Özdinler, P. H., & Macklis, J. D. (2006). IGF-I specifically enhances axon outgrowth of corticospinal motor neurons. Nature Neuroscience, 9(11), 1371–1381. https://doi.org/10.1038/nn1789

Preissl, S., Fang, R., Huang, H., Zhao, Y., Raviram, R., Gorkin, D. U., Zhang, Y., Sos, B. C., Afzal, V., Dickel, D. E., Kuan, S., Visel, A., Pennacchio, L. A., Zhang, K., & Ren, B. (2018). Single-nucleus analysis of accessible chromatin in developing mouse forebrain reveals cell-type-specific transcriptional regulation. Nature Neuroscience, 21(3), 432–439. https://doi.org/10.1038/s41593-018-0079-3

Rogge, G., Jones, D., Hubert, G. W., Lin, Y., & Kuhar, M. J. (2008). CART peptides: Regulators of body weight, reward and other functions. Nature Reviews Neuroscience, 9(10), 747–758. https://doi.org/10.1038/nrn2493

Rosenberg, A. B., Roco, C. M., Muscat, R. A., Kuchina, A., Sample, P., Yao, Z., Graybuck, L. T., Peeler, D. J., Mukherjee, S., Chen, W., Pun, S. H., Sellers, D. L., Tasic, B., & Seelig, G. (2018). Single-cell profiling of the developing mouse brain and spinal cord with split-pool barcoding. Science (New York, N.Y)., 360(6385), 176–182. https://doi.org/10.1126/science.aam8999

Ruder, L., & Arber, S. (2019). Brainstem Circuits Controlling Action Diversification. Annual Review of Neuroscience, 42(1), 485–504. https://doi.org/10.1146/annurev-neuro-070918-050201

Ruder, L., Schina, R., Kanodia, H., Valencia-Garcia, S., Pivetta, C., & Arber, S. (2021). A functional map for diverse forelimb actions within brainstem circuitry. Nature, 590(7846), 445–450. https://doi.org/10.1038/s41586-020-03080-z

Sahni, V., Engmann, A., Ozkan, A., & Macklis, J. D. (2020). Motor cortex connections. In Neural Circuit and Cognitive Development. Elsevier Inc. https://doi.org/10.1016/B978-0-12-814411-4.00008-1

Sahni, V., Itoh, Y., Shnider, S. J., & Macklis, J. D. (2021). Crim1 and Kelch-like 14 exert complementary dual-directional developmental control over segmentally specific corticospinal axon projection targeting. Cell Reports, 37(3), 109842. https://doi.org/10.1016/j.celrep.2021.109842

Sahni, V., Shnider, S. J., Jabaudon, D., Song, J. H. T., Itoh, Y., Greig, L. C., & Macklis, J. D. (2021). Corticospinal neuron subpopulation-specific developmental genes prospectively indicate mature segmentally specific axon projection targeting. Cell Reports, 37(3), 109843. https://doi.org/10.1016/j.celrep.2021.109843

Schieber, M. H. (2007). Chapter 2 Comparative anatomy and physiology of the corticospinal system. In A.A. Eisen & P. J. Shaw (Eds.), Handbook of Clinical Neurology (Vol. 82, pp. 15–37). Elsevier. https://doi.org/10.1016/S0072-9752(07)80005-4

Shim, S., Kwan, K. Y., Li, M., Lefebvre, V., & Šestan, N. (2012). Cis-regulatory control of corticospinal system development and evolution. Nature, 486(7401), 74–79. https://doi.org/10.1038/nature11094

Song, J. H., Choi, W., Song, Y.-H., Kim, J.-H., Jeong, D., Lee, S.-H., & Paik, S.-B. (2020). Precise Mapping of Single Neurons by Calibrated 3D Reconstruction of Brain Slices Reveals Topographic Projection in Mouse Visual Cortex. Cell Reports, 31(8), 107682. https://doi.org/10.1016/j.celrep.2020.107682

Svoboda, K., & Li, N. (2018). Neural mechanisms of movement planning: Motor cortex and beyond. Current Opinion in Neurobiology, 49, 33–41. https://doi.org/10.1016/j.conb.2017.10.023

Tasic, B. (2018). Single cell transcriptomics in neuroscience: Cell classification and beyond. Current Opinion in Neurobiology, 50, 242–249. https://doi.org/10.1016/j.conb.2018.04.021

Tasic, B., Menon, V., Nguyen, T. N., Kim, T. K., Jarsky, T., Yao, Z., Levi, B., Gray, L. T., Sorensen, S. A., Dolbeare, T., Bertagnolli, D., Goldy, J., Shapovalova, N., Parry, S., Lee, C., Smith, K., Bernard, A., Madisen, L., Sunkin, S. M., … Zeng, H. (2016). Adult mouse cortical cell taxonomy revealed by single cell transcriptomics. Nature Neuroscience, 19(2), 335–346. https://doi.org/10.1038/nn.4216

Tennant, K. A., Adkins, D. L., Donlan, N. A., Asay, A. L., Thomas, N., Kleim, J. A., & Jones, T. A. (2011). The organization of the forelimb representation of the C57BL/6 mouse motor cortex as defined by intracortical microstimulation and cytoarchitecture. Cerebral Cortex (New York, N.Y.: 1991), 21(4), 865–876. https://doi.org/10.1093/cercor/bhq159

Tomassy, G. S., De Leonibus, E., Jabaudon, D., Lodato, S., Alfano, C., Mele, A., Macklis, J. D., & Studer, M. (2010). Area-specific temporal control of corticospinal motor neuron differentiation by COUP-TFI. Proceedings of the National Academy of Sciences, 107(8), 3576–3581. https://doi.org/10.1073/pnas.0911792107

vsnnlab. (2021). AMaSiNe [MATLAB]. https://github.com/vsnnlab/AMaSiNe (Original work published 2020)

Woodworth, M. B., Greig, L. C., Kriegstein, A. R., & Macklis, J. D. (2012). SnapShot: Cortical Development. Cell, 151(4), 918-918.e1. https://doi.org/10.1016/j.cell.2012.10.004

Yao, Z., Liu, H., Xie, F., Fischer, S., Adkins, R. S., Aldridge, A. I., Ament, S. A., Bartlett, A., Behrens, M. M., Van den Berge, K., Bertagnolli, D., de Bézieux, H. R., Biancalani, T., Booeshaghi, A. S., Bravo, H. C., Casper, T., Colantuoni, C., Crabtree, J., Creasy, H., … Mukamel, E. A. (2021). A transcriptomic and epigenomic cell atlas of the mouse primary motor cortex. Nature, 598(7879), 103–110. https://doi.org/10.1038/s41586-021-03500-8

Zeisel, A., Muñoz-Manchado, A. B., Codeluppi, S., Lönnerberg, P., La Manno, G., Juréus, A., Marques, S., Munguba, H., He, L., Betsholtz, C., Rolny, C., Castelo-Branco, G., Hjerling-Leffler, J., & Linnarsson, S. (2015). Cell types in the mouse cortex and hippocampus revealed by single-cell RNA-seq. Science, 347(6226), 1138–1142. https://doi.org/10.1126/science.aaa1934

Zheng, G. X. Y., Terry, J. M., Belgrader, P., Ryvkin, P., Bent, Z. W., Wilson, R., Ziraldo, S. B., Wheeler, T. D., McDermott, G. P., Zhu, J., Gregory, M. T., Shuga, J., Montesclaros, L., Underwood, J. G., Masquelier, D. A., Nishimura, S. Y., Schnall-Levin, M., Wyatt, P. W., Hindson, C. M., … Bielas, J. H. (2017). Massively parallel digital transcriptional profiling of single cells. Nature Communications, 8(1), 14049. https://doi.org/10.1038/ncomms14049

